# Furanoditerpenoid biosynthesis in the bioenergy crop switchgrass is catalyzed by an alternate metabolic pathway

**DOI:** 10.1101/2021.03.30.437764

**Authors:** Andrew Muchlinski, Meirong Jia, Kira Tiedge, Jason S. Fell, Kyle A. Pelot, Lisl Chew, Danielle Davisson, Yuxuan Chen, Justin Siegel, John T. Lovell, Philipp Zerbe

**Author notes:** To whom correspondence should be addressed: Philipp Zerbe; ORCID ID: 0000-0001-5163-9523. These authors contributed equally to this work (Andrew Muchlinski, Meirong Jia, Kira Tiedge, Jason S. Fell). State Key Laboratory of Bioactive Substance and Function of Natural Medicines & NHC Key Laboratory of Biosynthesis of Natural Products, Institute of Materia Medica, Chinese Academy of Medical Sciences & Peking Union Medical College, Beijing 100050, China.

## Abstract

Specialized diterpenoid metabolites are important mediators of stress resilience in monocot crops. A deeper understanding of how species-specific diterpenoid-metabolic pathways and functions contribute to plant chemical defenses can enable crop improvement strategies. Here, we report the genomics-enabled discovery of five cytochrome P450 monooxygenases (CYP71Z25-29) that form previously unknown furanoditerpenoids in the monocot bioenergy crop switchgrass (*Panicum virgatum*). Combinatorial pathway reconstruction showed that CYP71Z25-29 catalyze furan ring addition to diterpene alcohol intermediates derived from distinct class II diterpene synthases, thus bypassing the canonical role of class I diterpene synthases in plant diterpenoid metabolism. Transcriptional co-expression patterns and presence of select diterpenoids in droughted switchgrass roots support possible roles of CYP71Z25-29 in abiotic stress responses. Integrating molecular dynamics, structural analysis, and targeted mutagenesis, identified active site determinants controlling distinct CYP71Z25-29 catalytic specificities and, combined with broad substrate promiscuity for native and non-native diterpenoids, highlights the potential of these P450s for natural product engineering.

**Significance Statement:** Diterpenoids play important roles in stress resilience and chemically mediated interactions in many plant species, including major food and bioenergy crops. Enzymes of the cytochrome P450 monooxygenase family catalyze the various functional decorations of core diterpene scaffolds that determine the large diversity of biologically active diterpenoids. This study describes the identification and mechanistic analysis of an unusual group of cytochrome P450 monooxygenases, CYP71Z25-29, from the bioenergy crop switchgrass (*Panicum virgatum*). These enzymes catalyze the furan ring addition directly to class II diterpene synthase products, thus bypassing the conserved pairwise reaction of class II and class I diterpene synthases in labdane diterpenoid metabolism. Insight into the distinct substrate-specificity of CYP71Z25-29 offers opportunity for engineering of furanoditerpenoid bioproducts.

## Introduction

Diverse networks of specialized metabolites impact plant fitness by mediating ecological interactions among plants, microbes and animals. Among these metabolites, diterpenoids play essential roles in plant defense and ecological adaptation (Tholl, 2015). For instance, chemically distinct diterpenoid blends confer pathogen and pest resistance in major global grain crops, including maize (*Zea mays*) and rice (*Oryza sativa*) (Schmelz et al., 2014; Murphy and Zerbe, 2020). Recent studies further suggest diterpenoid functions in mediating abiotic stress adaptation. For example, UV irradiation elicited diterpenoid accumulation and expression of the corresponding metabolic genes in rice (Park et al., 2013; Horie et al., 2015). Inducible diterpenoid formation was also observed in maize in response to oxidative, drought and salinity stress (Vaughan et al., 2014; Christensen et al., 2018; Mafu et al., 2018), and diterpenoid-deficient maize mutants show decreased resilience to abiotic perturbations (Vaughan et al., 2015). Expanding knowledge of diterpenoid diversity, associated metabolic pathways and functions across a broader range of monocot crop species can inform adaptive breeding and engineering strategies to improve crop environmental resilience (Bevan et al., 2017; Nelson et al., 2018; Bailey-Serres et al., 2019).

Switchgrass (*Panicum virgatum*) is a key species of the North American tallgrass prairie ecosystem and valued as a forage and biofuel crop for its high net energy yield and abiotic stress tolerance (Schmer et al., 2008; Liu et al., 2015; Lovell et al., 2021). Broad drought-induced alterations in carbohydrate, lipid, phenylpropanoid and terpenoid metabolism support a role of specialized metabolites in switchgrass abiotic stress resilience (Xingxing Li; Meyer et al., 2014; Pelot et al., 2018; Muchlinski et al., 2019).

Diterpenoids in monocot crops almost invariably belong to the group of labdane-type metabolites, and feature species-specific structures, bioactivities, and spatiotemporal regulation and distribution (Schmelz et al., 2014; Murphy and Zerbe, 2020). Diterpene synthases (diTPS) and cytochrome P450 monooxygenases (P450) are the key gatekeepers to diterpenoid diversity (Zerbe and Bohlmann, 2015; Banerjee and Hamberger, 2018). Rooted in the common C20 precursor, geranylgeranyl pyrophosphate (GGPP), the conserved pathway architecture en route to labdane diterpenoids recruits the combined activity of class II and class I diTPSs. After class II diTPS catalyzed conversion of GGPP into bicyclic prenyl pyrophosphate compounds of distinct stereochemistry and oxygenation, class I diTPSs facilitate the dephosphorylation and subsequent cyclization and/or rearrangement of these intermediates to generate various diterpenoid scaffolds (Peters, 2010). Functional decoration through the activity of P450s and other modifying enzyme classes then expands the structural complexity and bioactivity of plant diterpenoids (Banerjee and Hamberger, 2018). Over the past decades, numerous diterpenoid-metabolic diTPSs and P450s have been identified in maize, rice and wheat (*Triticum aestivum*) (reviewed in Schmelz et al., 2014; Murphy and Zerbe, 2020), and demonstrated that downstream of the central GGPP precursor, labdane diterpenoid biosynthesis is organized as modular metabolic networks, where pairwise reactions of functionally distinct enzymes create multiple pathway branches to readily increase product diversity (Xu et al., 2007; Morrone et al., 2011; Mafu et al., 2018; Murphy et al., 2018; Ding et al., 2019). By integrating genome-wide pathway discovery and combinatorial protein biochemical tools, our prior work identified a large and diverse diTPS family in switchgrass (*Panicum virgatum*) (Pelot et al., 2018) that yields an expansive diversity of diterpenoids, including several labdane-type compounds that occur, to current knowledge, uniquely in switchgrass (Pelot et al., 2018) (Fig. 1). Endogenous accumulation of several metabolites and expression of the corresponding biosynthetic genes in roots and leaves following abiotic stress support a role of terpenoids in switchgrass environmental adaptation (Pelot et al., 2018; Muchlinski et al., 2019); however, complete metabolic pathways, products, and their physiological functions remain to be resolved.

**Figure 1.**
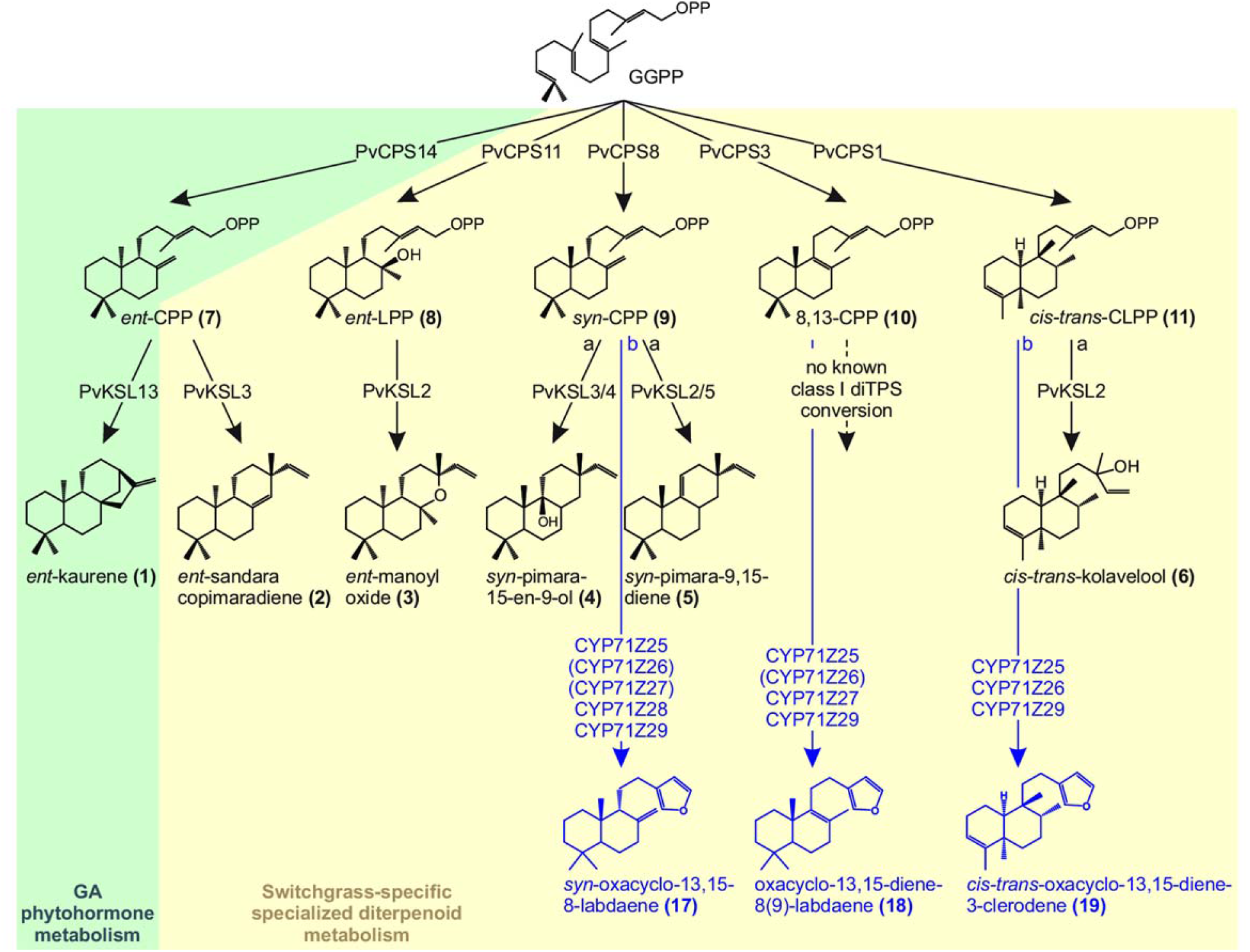
Switchgrass diterpenoid-metabolic network. Shown is an overview of identified biosynthetic pathways, involving monofunctional class II and class I diTPSs, as well as P450 enzymes (this study) that convert the central precursor GGPP into a range of diterpenoid metabolites with putative roles in general (gibberellin phytohormone; green) and specialized (yellow) diterpenoid metabolism.

Combining genomic studies, combinatorial enzyme assays, metabolite and transcript profiling, and protein structure-function studies revealed a group of five P450s of the CYP71 clan (CYP71Z25-29) that convert a range of diterpene scaffolds into furanoditerpenoid derivatives. P450-catalyzed addition of a furan ring directly to diterpene alcohol intermediates derived from class II diTPS activity illustrates a previously unrecognized alternative to the common labdane diterpenoid formation requiring pairwise class II and class I diTPS activity. Co-expression of functionally compatible *diTPSs* and *P450s* in switchgrass roots and drought-elicited accumulation of select diterpenoids support a role in abiotic stress responses. Mechanistic insight into CYP71Z25-29 catalysis enables resources for engineering a broad range of bioactive furanoditerpenoids.

## Results

### Identification of diterpenoid-metabolic P450s in the switchgrass genome

To elucidate P450 pathways for the functional decoration of the expansive spectrum of diterpenoid structures in switchgrass, we probed the genomic regions neighboring known switchgrass *diTPS* genes (*P. virgatum*; var. Alamo AP13; genome version v5.1) (Lovell et al., 2021, Pelot et al., 2018). A tandem pair of two *P450* genes, designated *CYP71Z26* (*Pavir.1KG382300*) and *CYP71Z27* (*Pavir.1KG382400*), co-localized on chromosome 1K in direct proximity to three class II *diTPS* genes, including the *cis-trans*-clerodienyl pyrophosphate (*cis-trans*-CLPP) synthase *PvCPS1* (*Pavir.1KG382200*) (12.5 kb and 43.5 kb, respectively), its paralog *PvCPS2* (*Pavir.1KG382115*), and an additional putative *diTPS* (*Pavir.1KG382110*) (Fig. 2A). An additional P450 candidate, *CYP71Z25* (*Pavir.1KG341400*), was identified distantly (1.3 Mb from *CYP71Z27*) on chromosome 1K. Orthology networks including the genomes of switchgrass, the diploid switchgrass relative *P. hallii* (DOE-JGI) (Lovell et al., 2018), and *Setaria italica* (Bennetzen et al., 2012) identified five additional switchgrass P450s, while only one in *S. italica* and two possible paralogs in *P. hallii* were observed (Fig. S1). Notably, the paralogs, *Pahal.A02218* and *Pahal.A02220*, were also clustered in the genome of *P. hallii* (var. filipes FIL2; version v2.0) (Lovell et al., 2018) and co-localized with two class II diTPSs, *Pahal.A02215* and *Pahal.A02217*, with predicted *cis-trans*-CLPP and 8,13-CPP synthase activities, respectively (Pelot et al., 2018) (Fig. 2B). Among the remaining identified switchgrass candidates, only two, *CYP71Z28* (*Pavir.1NG304500*) and *CYP71Z29* (*Pavir.1NG309700*), represented full-length open reading frames and are located distantly (~552 kb) from each other on chromosome 1N (Fig. 2A). CYP71Z25-29 showed high protein sequence identity (90-99%) among the five switchgrass proteins and 82-91% to the two *P. hallii* homologs (Fig. 2C, Fig. S1).

**Figure 2.**
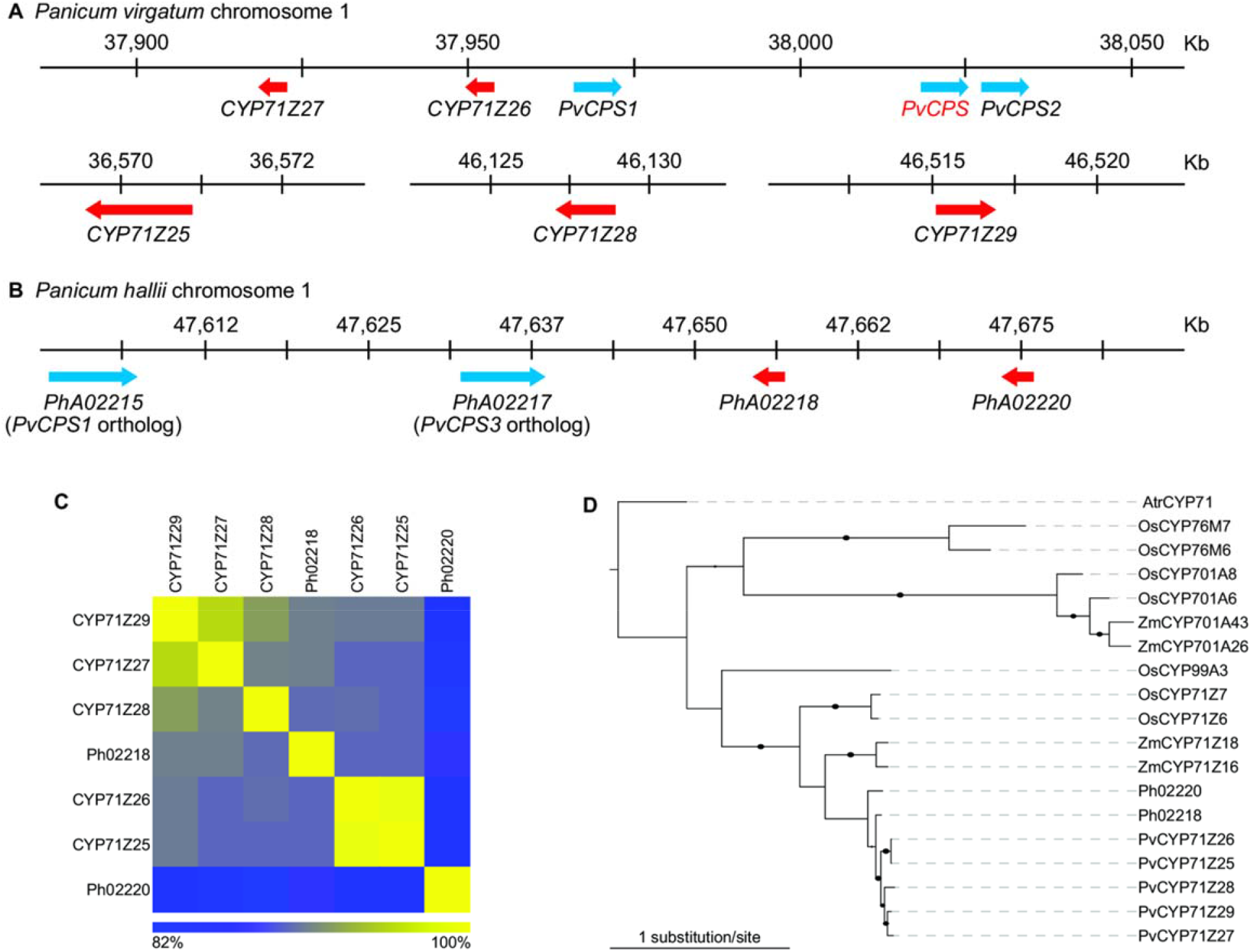
Discovery of switchgrass (Panicum virgatum) cytochrome P450 monooxygenases. **(A-B)** Genomic co-location of P450 genes (red) with class II diterpene synthases (blue) in the *P. virgatum* (var. Alamo AP13) genome (version v5.1) and the genome of the diploid switchgrass relative *P. hallii* (var. filipes FIL2, version v3.1) (DOE-JGI(b); Loveli et al. 2018). **(C)** Heat map illustrating the amino acid sequence identity of the five identified switchgrass P450 candidates, CYP71Z25-29, and orthologous P45Os from the diploid switchgrass relative *P. hallii*. **(D)** Maximum-likelihood phylogenetic tree of CYP71Z25-29 with predicted or characterized diterpenoid-metabolic P45Os from other monocot crops (Table S4). Tree rooted with the uncharacterized CYP71 from *Amborella trichopoda*. Branches with bootstrap support greater than 80% (500 repetitions) are depicted by black circles.

Phylogenetic analysis with characterized diterpenoid-metabolic P450s from related Poaceae species placed the switchgrass proteins in a distinct clade together with the *P. hallii* (Fig. 2D). This branch showed the closest relationship with members of the CYP71 clan with known roles in specialized diterpenoid metabolism in maize and rice (Wu et al., 2011; Mao et al., 2016; Mafu et al., 2018; Ding et al., 2019), thus indicating related, yet distinct functionalities.

### Switchgrass CYP71Z enzymes form distinct furanoditerpenoids

To functionally characterize the identified switchgrass P450s (CYP71Z25-29), we employed combinatorial pathway reconstruction assays of codon-optimized, N-terminally modified P450 constructs with functionally distinct diTPSs and a maize cytochrome P450 reductase (*Zm*CPR2) using an established *E. coli* expression platform (Morrone et al., 2010; Murphy et al., 2019). Following the typical pathway organization of plant labdane diterpenoid metabolism (Peters, 2010), we first tested the co-expression of each P450 with different pairs of class II and class I diTPSs that produce six core diterpenoid scaffolds formed by the switchgrass diTPS family (Pelot et al., 2018), namely *ent*-kaurene **1** (derived from *ent*-copalyl pyrophosphate, CPP **7**), *ent*-sandaracopimaradiene **2** (derived from **7**), *ent*-manoyl oxide **3** (derived from *ent*-labdadienol pyrophosphate, LPP **8**), *syn*-pimara-15-en-9-ol **4** (derived from *syn*-CPP **9**), *syn*-pimara-9,15-diene **5** (derived from **9**), and *cis-trans*-kolavelool **6** (derived from *cis-trans*-clerodienyl pyrophosphate, CLPP **11**). For compound numbering and structures see Fig. 1 & Fig. S2. When compared to the products of the combined class II and class I diTPS activity alone, trace amounts of P450 products were detected only for CYP71Z25, CYP71Z26, and CYP71Z29 when co-expressed with *Pv*CPS1 and *Pv*KSL2 that form **11** and **6**, respectively (Fig. 3A; Fig. S3). The respective P450 products featured near identical fragmentation patterns with characteristic mass ions of *m/z* 286, 191, 177, 95, and 81, indicative of an oxygenated labdane scaffold. Notably, presence of these P450 products when co-expressed with *Pv*CPS1 and *Pv*KSL2 coincided with this diTPS pair yielding the lowest product formation of all tested diTPS combinations, resulting in a higher accumulation of the dephosphorylated *cis-trans*-CLPP 15-hydroxy derivative *cis-trans*-kolavenol **16**, due to the activity of endogenous *E. coli* phosphatases.

**Figure 3.**
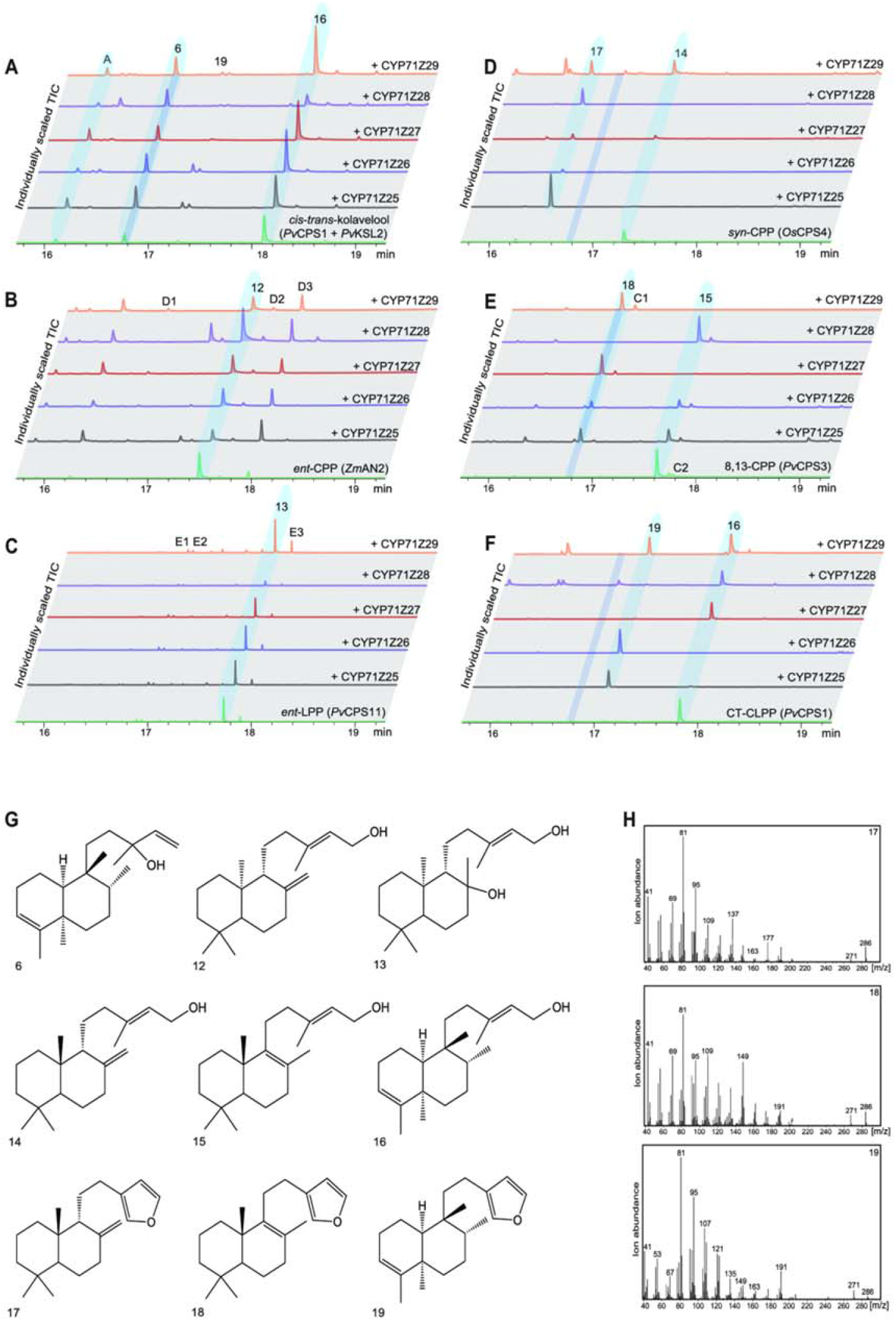
Functional characterization of switchgrass CYP71Z25-29. **(A-F)** Individually scaled total ion GC-MS chromatograms of enzyme products resulting from E. *coli* co-expression of each P450 candidate (CYP71Z25-29) with different diterpene synthases: **(A)** PvCPS1 and PvKSL2 producing the diterpenoid scaffold cis-trans-koiavelool **6**, **(B)** ZmAN2 producing *ent*-copalyl pyrophosphate (CPP) **7**, **(C)** PvCPS11 producing enf-labdadienyi pyrophosphate (LPP) **8**, **(D)** *Oryza saliva* CPS4 producing syn-CPP **9**. **(E)** PvCPS3 producing B.13-CPP **10**. and **(F)** PvCPS1 producing *cis-trans*-clerodienyl pyrophosphate (CT-CLPP) **11**. Compounds labeled as C1-2, D1-3 and E1-3 indicate unidentified terpenoid products based on mass spectral patterns. Unlabeled peaks represent non-terpenoid compounds derived from E. *coli* expression cultures. **(G)** Structures of relevant enzyme products. Structures of **17, 18**, and **19** were verified by NMR analysis and absolute stereochemistry assigned based on the corresponding class II diterpene synthase products. (H) Mass spectra of identified P450 furanoditerpenoid products.

This result, in conjunction with the genomic co-localization of several P450 candidates with *PvCPS1* or *PvCPS2*, supported the hypothesis that CYP71Z25-29 function in combination with only class II diTPSs. To test this hypothesis, *E. coli* co-expression assays were conducted pairing each P450 candidate with individual class II diTPSs producing the distinct prenyl pyrophosphate products known in switchgrass: **7**, **8**, **9**, **11** and 8,13-CPP **10** (Pelot et al., 2018) (Fig. S2). GC-MS analysis of the respective reaction products showed product formation by all P450s, although with apparent differences in substrate specificity (Fig. 3B-F). It should be noted that GC-MS analysis detected the dephosphorylated class II diTPS products due to the activity of endogenous *E. coli* phosphatases (*ent*-copalol **12**, *ent*-labdadienol **13**, *syn*-copalol **14**, 8,13-copalol **15**, and *cis-trans*-kolavenol **16**. No P450 products were observed with **12** or **13** as substrates (Fig. 3B-C). Conversion of **16** was observed for CYP71Z25, CYP71Z26 and CYP71Z29, whereas CYP71Z27 and CYP71Z28 showed no activity. Importantly, the resulting P450 product featured the same retention time and a near-identical fragmentation pattern as compared to the product observed when combing the *cis-trans*-CLPP (**11**) synthase *Pv*CPS1 with the *cis-trans*-kolavelool **6** synthase *Pv*KSL2 and a P450 candidate (Fig. 3A,F). Conversion of **15** was observed for CYP71Z27, CYP71Z29, and partially for CYP71Z25 and CYP71Z26, whereas CYP71Z28 was largely inactive with this substrate (Fig. 3E). By contrast, CYP71Z28 showed high activity with **14** as a substrate, as did CYP71Z25 and CYP71Z26, whereas CYP71Z27 and CYP71Z29 showed only incomplete substrate conversion (Fig. 3D). For all observed P450 products, fragmentation patterns featured *m/z* 286, 191, and 177 mass ions consistent with oxygenated labdane diterpenoid structures (Fig. 3G-H).

To determine the structure of selected P450 products, enzymatically produced and purified compounds were subject to 1D and 2D NMR (HSQC, COSY, HMBC and NOESY) analysis and compared to previously reported NMR data where available. This approach revealed a shared structural scaffold of the P450 products that contains the individual class II diTPS-derived diterpene backbones with a furan ring addition at the C15-C16 position (Fig. 3G). Furanoditerpenoid products identified here included *syn*-15,16-epoxy-8(17),13(16),14-triene **17** derived from the *Pv*CPS8 product **14** (Fig. S4), 15,16-epoxy-8,13(16),14-triene **18** derived from the *Pv*CPS3 product **15** (Fig. S5), and *cis-trans*-15,16-epoxy-cleroda-3,13(16),14-triene **19** derived from the *Pv*CPS1 product **16** (Fig. S6). The stereochemistry of the P450 products was assigned on the basis of the respective class II diTPS products (Pelot et al., 2018).

Considering the substrate promiscuity of CYP71Z25-29, we next investigated the capacity of these P450s to convert alternate class II diTPS products. Indeed, with the exception of CYP71Z28, all P450s formed (+)-15,16epoxy-8(17),13(16),14-triene **26** when co-expressed with a diTPS producing (+)-CPP **20** ((+)-copalol **23**), a class II diTPS product not currently known in switchgrass, but formed by diTPSs of other monocot crops such as wheat and maize (Wu et al., 2012; Murphy et al., 2018) (Figs. S7 and S8). In addition, CYP71Z25 and CYP71Z26 showed, albeit low, activity when co-expressed with the *trans-cis*-neo-clerodienyl pyrophosphate (*trans-cis*-CLPP **22**; *trans-cis*-kolavenol **25**) synthase of *Salvia divinorum, SdCPS2* (Pelot et al., 2017), forming the core precursor to salvinorin A, a neoclerodane furanoditerpenoid with potential use for the treatment of drug addiction and neuropsychiatric disorders (Kivell et al., 2014). The two major P450 products showed signature mass ions of *m/z* 286 or *m/z* 288, indicating a **25**-derived furanoditerpenoid **28** and *trans-cis*-cleroda-3,12-dien-15,16-diol **29** structures, respectively (Fig. S7).

Formation of furanoditerpenoids through the coupled activity of the focal P450s and functionally distinct class II diTPSs warranted a deeper investigation of the nature of the P450 substrate. While class II diTPSs form prenyl pyrophosphate products with a characteristic pyrophosphate group at C15 (Peters, 2010), as mentioned above, expression of class II diTPSs in heterologous plant or microbial host systems typically yields the corresponding C15-hydroxy derivatives formed through the activity of endogenous phosphatases (Mafu et al., 2018; Pelot et al., 2018; Ding et al., 2019). To clarify the catalytic preference of CYP71Z25-29, substrate feeding experiments were conducted by adding enzymatically produced and purified compounds of **15** or **16** to *E. coli* cultures expressing CYP71Z25 or CYP71Z29. For both tested P450-substrate combinations, conversion of the 15-hydroxy diterpene substrates into the respective furanoditerpenoids was observed, whereas no conversion was detected in control samples expressing plasmids carrying *Zm*CPR2 alone (Fig. S9). Substantial limitations in purifying the corresponding prenyl pyrophosphate compounds prevented the possibility to conduct complementary feeding assays within the scope of this study.

### Structure-guided mutagenesis identifies active site determinants of P450 catalytic specificity

To investigate the catalytic mechanism underlying the unusual activity of CYP71Z25-29, homology models were generated for each P450 using the recently reported crystal structure of *Salvia miltiorrhiza* CYP76AH1 (Gu et al., 2019) as a template. Given a protein identity of only ~ 42% to the target P450s (Fig. S10), an iterative homology modeling and energy minimization approach employing relaxed template protein structures (Nivón et al., 2013; Pei and Grishin, 2014) was used to generate high-quality models. The resulting lowest energy model was used for ligand docking of the heme co-factor into the individual active sites. The generated structural models showed root-mean-square deviation (RMSD) values of 0.67-0.75 as compared to the template, thus representing high-quality reproduction of common secondary structures and placement of active site residues as demonstrated, for example, for the heme-anchoring cysteine C421 (Fig. S10). The five diterpenoid substrates tested in this study were then docked into the heme-model complex with three conceivable substitution arrangements: 15-hydroxy and 15-pyrophosphate structures as alternate substrates and 15,16-dihydroxy derivatives as intermediate or product of the P450-catalyzed oxygenation reaction (Figs. S2 and S10). Comparison of the interaction energy (IE) of all generated substrate-P450 docking poses showed that the average IE was most favorable for the 15-hydroxy and 15-16-dihydroxy substrates, whereas the 15-pyrophosphate structure was energetically less favorable (Table S1). The difference in IE between the 15-hydroxy and 15-pyrophosphate ligands can presumably be attributed to the far larger volume and electrostatic charge of the pyrophosphate moiety.

The various modeled substrate-protein complexes were then used to investigate active site residues with possible impact on the distinct substrate specificity of CYP71Z25-29. A total of 15 active site residues associated with the six known CYP71 substrate recognition sites (SRS) (Dueholm et al., 2015) were identified that were located proximal to the docked substrates and showed residue variation among CYP71Z25-29 and related members of the CYP71Z subfamily (Fig. 4A). To investigate the catalytic impact of these residues, protein variants of CYP71Z25 and CYP71Z27, as the functionally most contrasting P450s, were generated via site-directed mutagenesis and functionally characterized by *E. coli* co-expression with individual class II diTPSs producing native (**14, 15, 16**) or non-native (**23, 24, 25**) P450 substrates. Reciprocal mutagenesis of all 15 identified residues between CYP71Z25 and CYP71Z27 (F/Y81, F/S86, V/I89, N/D95, S/T187, L/Q188, A/G215, V/Y218, R/Q220, V/L226, E/D283, T/I288, L/M293, S/T346, M/I463) resulted in a near-complete loss of function of the corresponding CYP71Z25 variant (Fig. 4B, Fig. S10). By contrast, the reciprocal multi-residue variant of CYP71Z27 featured an altered product profile largely reflecting the wild type products of CYP71Z25.

**Figure 4.**
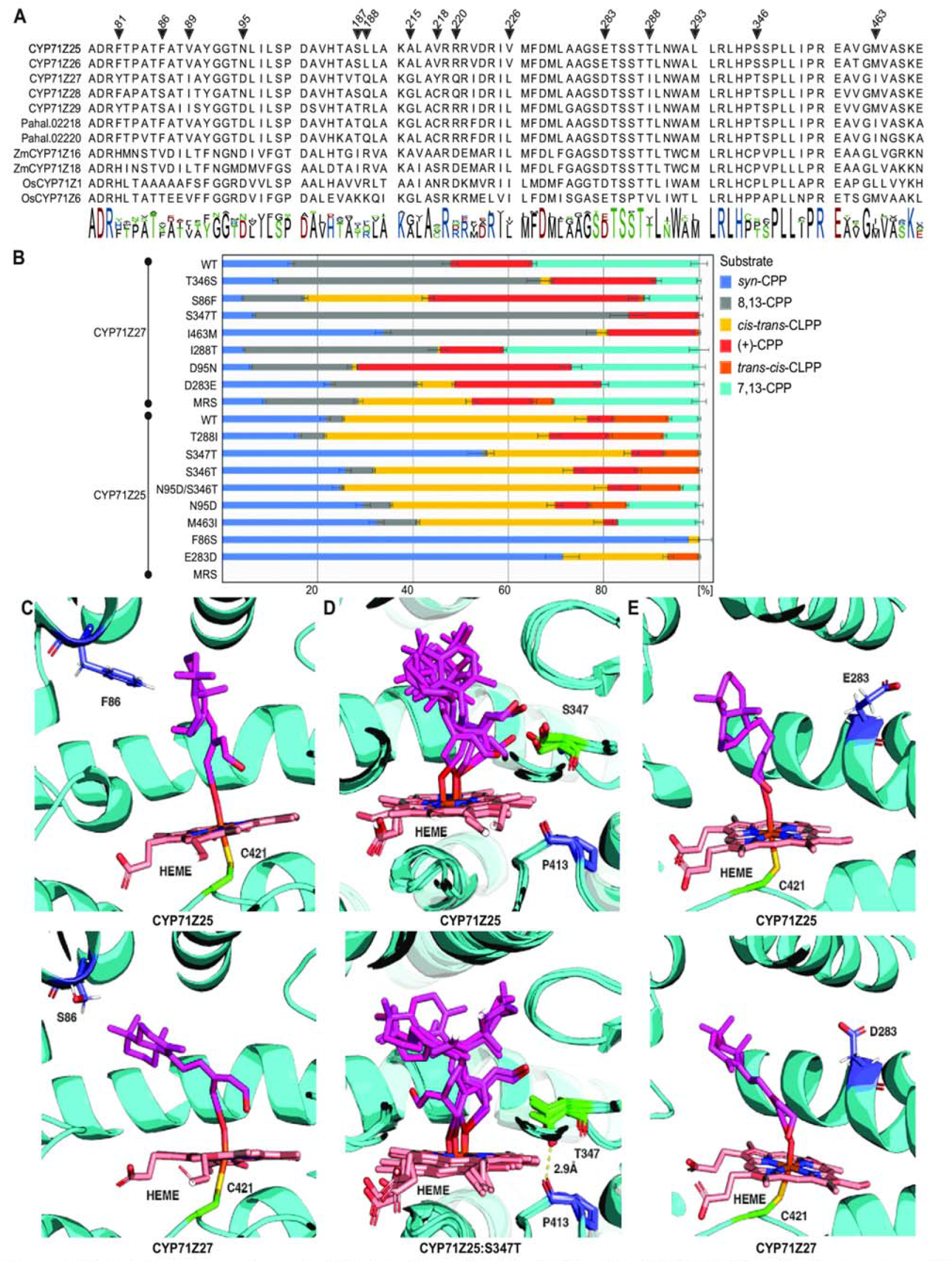
**(A)** Protein sequence alignment of Substrate Recognition Site (SRS) motifs of CYP71Z25-29 and related members of the CYP71Z subfamily of known or predicted function. **(B)** Analysis of standardized substrate conversion by CYP71Z25 and CYP71Z27 SDM variants. *E. coli* co-expressions for each variant with native (14, 15, 16) and non-native (23, 24, 25) switchgrass substrates were earned out in triplicate and normalized to the internal standard 1-eicosane and the OD600 at time of induction of protein expression. Error bars indicate standard deviation of replicates from the mean. **(C)** Active sites of CYP71Z25 (right) and CYP71Z27 (left) with heme (magenta), substrate (pink), and residues C412 (green) and S/F86 (blue). **(D)** Structural overlay of the five lowest IE active sites of CYP71Z25 (left) and CYP71Z25-S347T variant (right) with 15 (8,13-copalol) substrate (pink) and heme (magenta) docked. Residues S/T347 (green), P413 (blue), and corresponding hydrogen bonding (yellow dashes) with distance in Angstrom are highlighted. **(E)** Active sites of CYP71Z25 (right) and CYP71Z27 (left) with heme (magenta), substrate (pink), and residues C412 (green) and E/D283 (blue).

Strikingly, the single residue variant CYP71Z27:S86F showed a product profile similar to that observed for the multi-residue variant of CYP71Z27, converting all six tested diterpene alcohol substrates (Fig. 4B,C). By contrast, the reciprocal CYP71Z25:F86S variant showed only trace product amounts, again comparable to the multi-residue variant of CYP71Z25. Most notably, this protein variant showed substantial activity in producing **19** making up 23.7±0.6% of the product profile. Analysis of additional single residue variants revealed S347 hydrogen bonds with the docked hydroxy substrates and is conserved in CYP71Z25-29 (Fig. 4A,D). Substitution of S347 for a Thr impaired enzyme activity in CYP71Z27 with the exception of the conversion of 8,13-CPP-derived diterpene alcohols, whereas the same mutation had a lesser impact on CYP71Z25 catalysis reducing predominantly the conversion of **15** by 2.6% and **24** by 6.4% (Fig. 4B). Consistent with the observed protein variant activities, docking of the **15**-derived substrate into the active site crevice of CYP71Z25 and the corresponding S347T variant suggests that exchange of the native S347 residue to a Thr results in hydrogen bond formation with a neighboring Pro residue (P413), rather than the hydrogen bond formed with the substrate in the wild type enzymes (Fig. 4D). Reciprocal exchange of residues E/D283 significantly reduced product formation in CYP71Z25, especially with regards to the conversion of **15, 23, 24** (Fig. 4B,E), whereas the corresponding variant CYP71Z27:D283E showed only minor product changes with **19** as an additional product contributing 7.7±0.1% to the profile.

Mutagenesis of the remaining selected residues in CYP71Z25 and CYP71Z27, namely N/D95, T/I288, S/T346 and M/I463, showed limited impact on P450 catalytic specificity, including the formation of **19** as a product absent in the wild type enzyme by the CYP71Z27:D95N and CYP71Z27:T346S variants. Furthermore, the CYP71Z25:T288I variant and an additional double mutation, CYP71Z25:N95D/S346T showed a 9% increase in producing **28** when co-expressed with the *trans-cis*-CLPP synthase *Sd*CPS2 (Fig. S4).

### P450-derived furanoditerpenoids show drought-elicited formation in switchgrass roots

Previous work demonstrated that select diTPS transcripts and corresponding enzyme products, including **3** and **4**, accumulate in switchgrass (Alamo) leaves and roots exposed to below-ground oxidative stress (Pelot et al., 2018). To determine if furanoditerpenoid biosynthesis follows similar patterns, we examined P450-derived furanoditerpenoids in switchgrass plants exposed to four weeks of drought stress. Targeted GC-MS metabolite profiling of organic solvent extracts of droughted switchgrass roots and well-watered control plants illustrated an, albeit moderate, accumulation of several diterpenoid compounds, including **4**, a yet unidentified diterpenoid, and two metabolites, **30** and **31**, that show mass fragmentation patterns significantly similar but distinct from the identified furanoditerpenoid P450 products, suggesting that these compounds may represent further functionalized derivatives (Fig. 5A&B, Fig. S11). Specifically, mass ions of *m/z* 177 and *m/z* 192 indicated a *syn*-labdane backbone of compound **30**. Compound **31** featured a prominent *m*/*z* 148 possibly indicating an 8,13-labdane scaffold, as well as a *m/z* 304 mass fragment suggesting that this compound carries an additional hydroxy group (Fig. 5B). Low abundance of the identified diterpenoids prevented their purification from plant material for structural verification. Notably, all identified diterpenoids were present in both droughted and control plants with only compound **31** showing a moderate accumulation during the course of the drought treatment (Fig. S11).

**Figure 5.**
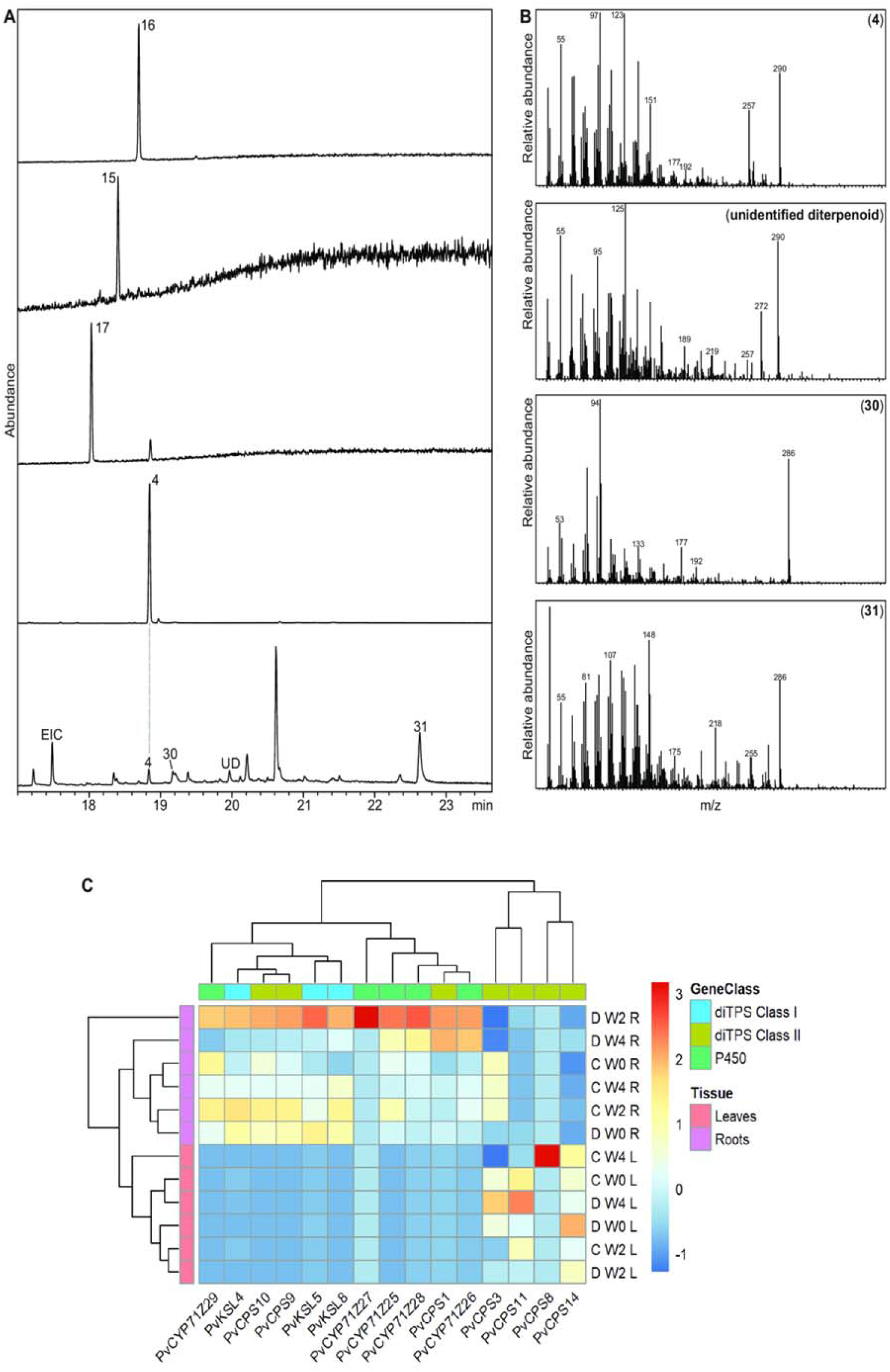
GC-MS total ion chromatograms **(A)** and mass spectra **(B)** of diterpenoîds detected in organic solvent extracts of switchgrass roots identified diterpenoids 4, an unidentified diterpenoid (UD) and predicted furanoditerpenoids (compounds 30 and 31) as possible derivatives of products of CYP71Z25-29. **(C)** Hierarchical cluster analysis of select diTPS and CYP71Z25-29 gene expression profiles from drought stressed tissues. Samples were collected before starting treatment (WO), two weeks (W2), and four weeks (W4) of drought stress treatment. L=leaves, R=roots, Drought stressed=D, well-watered=C.

Additional differential gene expression analysis using the same tissue samples was used to assess *diTPS* and *P450* gene expression patterns in droughted and control leaf and root tissue (Fig. 5B). Gene expression of *CYP71Z25-29* and the majority of diterpenoid-metabolic diTPSs, including the *cis-trans*-CLPP synthase *PvCPS1*, the predicted *syn*-CPP synthases *PvCPS9* and *PvCPS10* and the downstream-acting class I *diTPS PvKSL4, PvKSL5*, and *PvKSL8*, was detected only in roots consistent with the presence of compound 4 and predicted furanoditerpenoids. By contrast, the 8,13-CPP (10) synthase *PvCPS3* was moderately expressed in both organs, whereas the *ent*-LPP (8) synthase *PvCPS11* and the putatively gibberellin-biosynthetic *ent*-CPP (7) synthase *PvCPS14* were expressed only in leaves. Distinct from its homologs, *PvCPS9* and *PvCPS10*, the *syn*-CPP (9) synthase *PvCPS8* was expressed only in leaves after four weeks of treatment in well-watered plants, suggesting an expression profile at only later developmental stages. Drought-induced gene expression of root-specific specialized *diTPSs* and *CYP71Z25-29* was observed to be moderate with an increase until two weeks of drought treatment, followed by a decreased transcript abundance at four weeks of treatment (Fig. 5B). In leaves, expression of the specialized class II *diTPSs PvCPS3* and *PvCPS11* was highest after four weeks of drought treatment, whereas *PvCPS8* showed no drought-elicited transcript accumulation.

## Discussion

Crop improvement strategies increasingly benefit from knowledge of gene-to-metabolite relationships that contribute to desired crop traits and serve as resources for molecular crop engineering or breeding (Jez et al., 2016). Particularly, knowledge of the dynamic networks of plant specialized metabolites that enable plants to adapt to environmental pressures is needed in light of exacerbating crop losses caused by climate shifts and associated pest and disease damage (Savary et al., 2019). Diterpenoids serve as key components of biotic and abiotic stress resilience in rice and maize (Schmelz et al., 2014; Murphy and Zerbe, 2020), and stress-inducible, species-specific diterpenoid networks have also been discovered in other food and bioenergy crops such as switchgrass, foxtail millet and wheat, although their physiological functions are less well understood (Wu et al., 2012; Zhou et al., 2012; Schmelz et al., 2014; Pelot et al., 2018; Ding et al., 2019; Karunanithi et al., 2020). Prior studies identified an expansive *diTPS* gene family in allotetraploid switchgrass that forms specialized diterpenoids both common among the grass family and, to current knowledge, uniquely present in switchgrass (Pelot et al., 2018). The discovery of a group of functional *P450* genes, *CYP71Z25-29*, that function as furanoditerpenoid synthases via an alternate pathway independent of class I diTPS activity provides a deeper understanding of the divergence of switchgrass diterpenoid metabolism and how the natural modularity of diterpenoid-biosynthetic pathways drives the evolution of complex, lineagespecific blends of bioactive metabolites.

Identification of eight *CYP71Z-type* genes in switchgrass, as compared to only one paralog in *S. italica* and two genes in the diploid switchgrass relative *P. hallii*, suggests an expansion of this P450 group after the split from *P. hallii* approximately 8 MYA (Lovell et al., 2021). A close phylogenetic relationship among the functional members, CYP71Z25-29, indicates a shared evolutionary origin preceding gene duplication and sub/neo-functionalization events common in diterpenoid pathway evolution (Zi et al., 2014). Localization of *CYP71Z26* and *CYP71Z27* on chromosome 1K, along with a near-identical gene arrangement of two paralogs in *P. hallii*, whereas *CYP71Z25, CYP71Z28* and *CYP71Z29* are located on subgenome K, suggests that gene family expansion was associated with switchgrass subgenome expansion approximately 4.6 MYA (Lovell et al., 2021).

Diterpenoid-forming members of the CYP71Z subfamily also exist in maize and rice, where they catalyze position-specific hydroxylation, carboxylation or epoxidation reactions in various labdane scaffolds produced by the pairwise activity of class II and class I diTPS enzymes (Wu et al., 2011; Mafu et al., 2018; Ding et al., 2019). Consistent with their phylogenetic distance from maize and rice CYP71Z enzymes, functional characterization of switchgrass CYP71Z25-29 demonstrated a rare P450 activity in catalyzing the addition of a furan ring at C15-C16 of a range of different labdane scaffolds. The only known example of P450-mediated furan ring formation is the biosynthesis of the monoterpenoid menthofuran catalyzed by CYP71A32 in members of the *Mentha* genus (Bertea et al., 2001). Strikingly, contrasting known labdane diterpenoid pathways that invariably require the activity of class II and class I diTPSs, CYP71Z25-29 showed no detectable activity in converting class I diTPS products. Instead, efficient furanoditerpenoid synthase activity of CYP71Z25-29 was observed in co-expression assays with class II diTPSs alone (Fig. 3). Feeding assays with 15-hydroxy derivatives of select class II diTPS products illustrate that these diterpene alcohols rather than the 15-pyrophosphate compounds are the preferred CYP71Z25-29 substrates. This substrate preference is further supported by molecular ligand docking studies showing favorable interaction energies for 15-hydroxy and 15,16-dihydroxy intermediates as compared to the corresponding pyrophosphate structures. The observed furanoditerpenoid synthase activity of CYP71Z25-29 supports the presence of an alternative labdane diterpenoid pathway route in switchgrass that bypasses the common conversion of class II diTPS products by class I diTPSs and instead involves phosphatase-mediated dephosphorylation and subsequent CYP71Z25-29-catalyzed conversion of the resulting diterpene alcohol intermediates (Fig. 6A). Based on the conversion of the 15-hydroxy labdane substrates by CYP71Z25-29, and similar interaction energies observed for 15-hydroxy and 15,16 dihydroxy labdane intermediates in molecular docking analyses, a catalytic mechanism can be proposed that proceeds through initial substrate hydroxylation at C-16 and subsequent deprotonation and ring closure at C-15, followed by hydride shifts to facilitate dehydration and formation of the C13-C16 and C14-C15 double bonds (Fig. 6B). Genomic clustering of *CYP71Z25* and *CYP71Z26/27* with the class II diTPSs *PvCPS2* and *PvCPS1*, respectively (Fig. 1A), and co-expression of furanoditerpenoid synthase genes with several class II *diTPS* (including *PvCPS1*) in switchgrass roots (Fig. 5B) further support this hypothesis. While no terpenoid-metabolic plant phosphatases have yet been described, 15-hydroxy derivatives of class II diTPS products are commonly observed in co-expression assays using microbial or *Nicotiana benthamiana* platforms (Morrone et al., 2010; Mafu et al., 2018; Pelot et al., 2018; Ding et al., 2019). Indeed, at least ten predicted acid phosphatase and lipid phosphate phosphatase genes are present in the switchgrass genome (var. Alamo AP13; version v5.1) (Lovell et al., 2021) that are located across chromosomes 1N and 1K.

**Figure 6.**
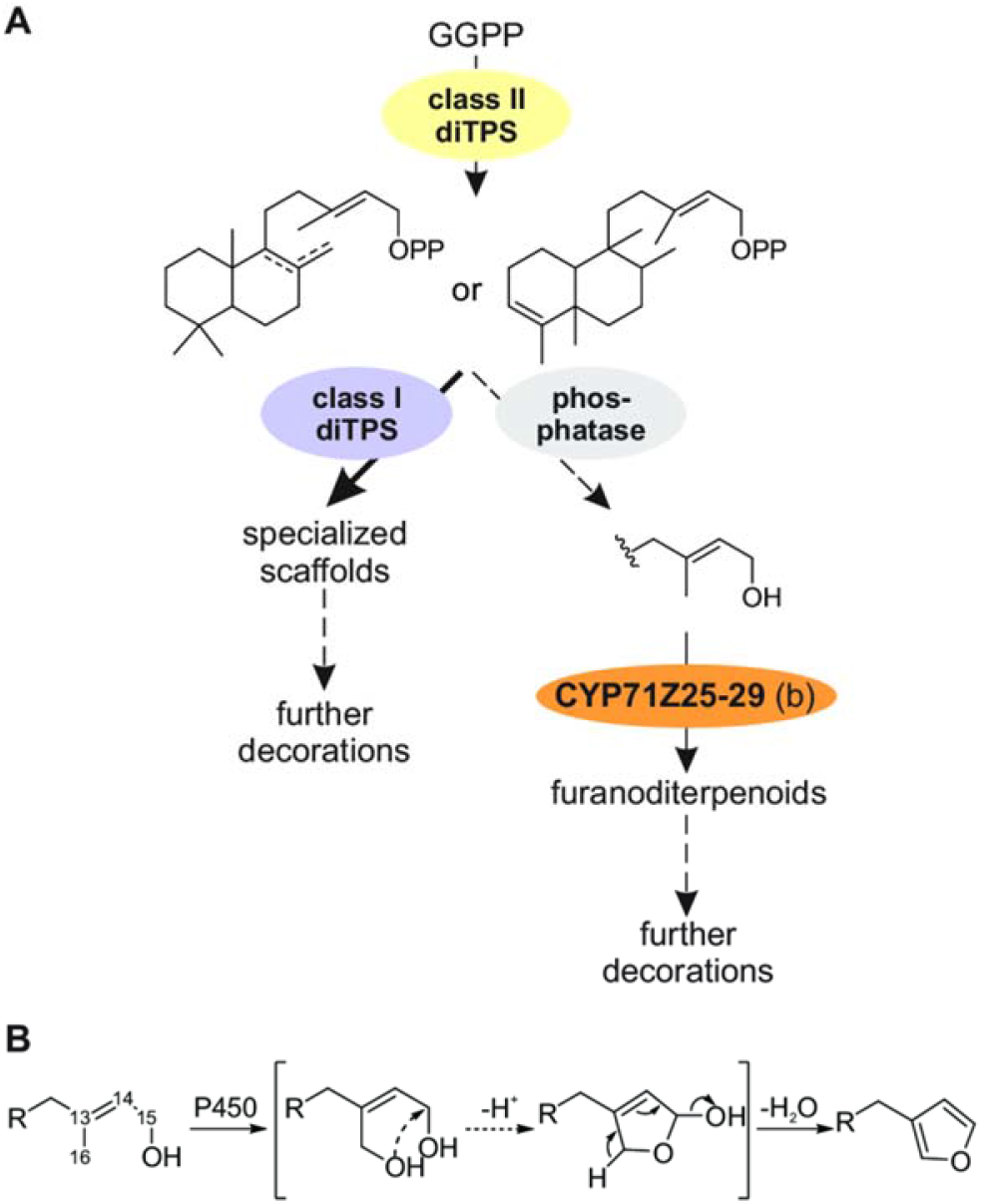
**(A)** Scheme of the postulated formation of diterpenoids in switchgrass. Derived from the common precursor geranylgeranyl pyrophosphate (GGPP) class II diterpene synthase products of different structure and stereochemistry can be formed either by further conversion through class I diterpene synthases and additional possible functional modifications or via cleavage of the pyrophosphate group by yet unidentified phosphatases and conversion of the resulting alcohol intermediates by CYP71Z25-29 to yield different furanoditerpenoids structures. **(B)** Proposed mechanism of furan ring addition catalyzed by CYP71Z25-29 that proceeds through oxidation of the C16 methyl group to form a dihydroxy intermediate, ring closure via deprotonation, and furan ring formation by dehydration.

The primary formation of class I diTPS products rather than furanoditerpenoids when coexpressing class II diTPSs, class I diTPSs and CYP71Z25-29 may suggest that the combined activity of class II and class I diTPSs represents the dominant pathway route. Compartmentalization of plastidial class II and class I diTPSs and common P450 localization at the endoplasmic reticulum supports pairwise diTPS activity as the primary pathway. However, presence of putative furanoditerpenoids derived from the identified P450 products alongside the class I diTPS product **4** and other yet unidentified diterpenoids in switchgrass roots supports the co-occurrence of both pathway branches *in planta*. Co-expression patterns of select biosynthetic genes, including *CYP71Z25/26* as well as *PvKSL4* and *PvKSL5*, relevant for the formation of Syn-CPP-derived diterpenoids in switchgrass roots is consistent with this hypothesis. However, expression of the *syn*-CPP synthase *PvCPS8* only in leaves, may indicate that the predicted functional homologs *PvCPS9* and/or *PvCPS10* are more likely to serve in the biosynthesis of **17** and **4** in roots (Fig. 5B). Given the large size of the switchgrass diTPS family and the catalytic overlap of CYP71Z25-29, future biochemical and genetic studies will be required to precisely decode the interactions governing the modular formation of switchgrass-specific diterpenoids. Interestingly, the lack of CYP71Z25-29 activity with substrates of *ent*-stereochemistry (including the GA precursor *ent*-CPP) demonstrates a dedicated role of this P450 group in specialized metabolism and highlights a biochemical separation of furanoditerpenoid and general diterpenoid metabolism in switchgrass. Similar impacts of substrate-specificity on partitioning different diterpenoid branches have recently been described in maize, where specificity of two class I diTPSs, *Zm*KSL2 and *Zm*KSL4, for producing distinct positional isomers of *ent*-kaurene contributes to the partitioning of pathways toward dolabralexin and kauralexin diterpenoids (Ding et al., 2019).

Presence of furanoditerpenoids and other specialized diterpenoids (Fig. 5) (Pelot et al., 2018) in switchgrass roots may indicate a role in abiotic stress responses. Albeit at moderate levels, drought-induced gene expression of *CYP71Z25-28* and select *class II diTPS* in roots further supports the association of furanoditerpenoid biosynthesis and switchgrass drought stress responses. Stress-elicited accumulation of diterpenoids with allelopathic and anti-microbial bioactivities in rice roots are well-established (Schmelz et al., 2014), and recent maize studies also demonstrated the root accumulation of diterpenoid dolabralexins and sesquiterpenoid zealexins in response to drought, oxidative and salinity stress (Schmelz et al., 2014; Vaughan et al., 2015; Christensen et al., 2018; Mafu et al., 2018; Murphy and Zerbe, 2020). However, the metabolite abundances detected in switchgrass are lower as compared to stress-elicited terpenoid accumulation in rice and maize. Given the structural distinctiveness of switchgrass diterpenoids, it is conceivable that switchgrass furanoditerpenoids exert different bioactivities in switchgrass interorganismal and environmental interactions. It also appears likely that the detected switchgrass furanoditerpenoids represent lower abundant pathway intermediates that undergo further functional decorations to generate bioactive pathway end products. Indeed, among the approximately 400 known furanoditerpenoids broadly distributed across the plant kingdom, the vast majority feature extensive modifications of both the furan ring and the diterpene skeleton that may include variations of hydroxylation, carboxylation, lactonization, glycosylation and other transformations (Bao et al., 2016).

Integrating sequence comparison, molecular dynamics analysis and site-directed mutagenesis proved a powerful tool for identifying active site residues contributing to CYP71Z25-29 catalysis and substrate specificity. The relative ease by which the substrate specificity of CYP71Z25 and CYP71Z27 for labdane intermediates of different stereochemistry and double bond configuration could be altered through minor active site modifications, supports a rapid P450 functional divergence during the evolution of switchgrass diterpenoid metabolism. This is exemplified by a near-complete functional conversion of CYP71Z27 to the product profile of CYP71Z25 with a single S86 to F substitution. However, loss of activity in the reciprocal CYP71Z25 variant suggests that different active site positions play major roles in product specificity among the identified P450s. Indeed, proximity of the F86 aromatic ring to the docked substrate in CYP71Z27 supports a role in substrate orientation in the active site crevice, whereas introduction of a bulky Phe side chain in the CYP71Z25 cavity may lead to steric hindrance of substrate binding and catalysis (Fig. 3C). Further mechanistic insight was gained from substitution of a conserved Ser in the SRS5 domain shown to be imperative for catalysis by hydrogen-bonding to the substrate. Molecular dynamics results suggest that substitution of this position for Thr leads to reduced conformational flexibility and associated side chain rotations that are critical for hydrogen bond formation with the substrate. Paired with the substrate promiscuity of CYP71Z25-29, these mechanistic insights provide a foundation for producing a broader range of furanoditerpenoid natural products. Notably, many plant furanoditerpenoids have been associated with therapeutic bioactivities ranging from anti-allergic and anti-diabetic to anti-cancer and anti-viral efficacies, whereby the furan group serves as a key pharmacophore (Bao et al., 2016). In this context, the potential of combinatorial pathway reconstruction for diterpenoid manufacturing is highlighted by the capacity of CYP71Z25 and CYP71Z26 to form the corresponding di-hydroxy and furan-derivatives of **25**, the precursor in the biosynthesis of salvinorin A (Pelot et al., 2017), a natural product of *S. divinorum* that was identified as a drug candidate for treatment of drug addictions due to its agonistic activity on brain kappa-opioid receptors (Kivell et al., 2014). Increased catalytic specificity observed in two CYP71Z25 variants further underscores the potential of structure-guided protein engineering for enabling desired P450 activities as more mechanistic insight into diTPS and P450 functions is gained.

## Materials and Methods

### Gene synthesis

CYP71Z25-29 were synthesized as codon-optimized and N-terminally modified genes by replacement of the sequence upstream of the LPP motif with the leader sequence MAKKTSSKGK (Swaminathan et al., 2009) (Table S2) and individually inserted into MCS2 of a pETDuet-1 vector (www.emdmillipore.com) carrying the full-length, codon-optimized maize cytochrome P450 reductase (*Zm*CPR2) in MCS1. Gene synthesis and cloning were performed by DOE Joint Genome Institute (JGI) with support through a DNA Synthesis Award (#2568). In addition, the multi residue variants of CYP71Z25 and CYP71Z27 were obtained from GenScript (www.genscript.com).

### *E. coli* co-expression assays

Co-expression of diTPSs and P450s was conducted in an *E. coli* platform engineered for diterpenoid production (Morrone et al., 2010; Murphy et al., 2019). In brief, pIRS and pGGxC plasmids (Morrone et al., 2010) carrying class II diTPSs with distinct products [*ent*-CPP **7** *(Z. mays* AN2; (Harris et al., 2005) used here in place of the native *Pv*CPS14 due to its higher catalytic activity), *syn*-CPP **9** (*O. sativa* CPS4; (Xu et al., 2004) used here in place of the native *Pv*CPS8 due to its higher catalytic activity), (+)-CPP **20** (*Abies grandis* abietadiene synthase variant D621A; (Morrone et al., 2010)), 8,13-CPP **10** (*Pv*CPS3), 7,13-CPP **21** *(Grindelia robusta* 7,13-CPP synthase (Zerbe et al., 2015)), *ent*-LPP **8** (*Pv*CPS11), *cis-trans*-CLPP **11** (*Pv*CPS1), *trans-cis*-CLPP **22** (*Salvia divinorum* CPS2; (Pelot et al., 2016) were co-expressed with different P450 genes. For additional co-expression of class I diTPSs, respective genes sub-cloned into the pET28b(+) expression vector were used (Pelot et al., 2018). Constructs were co-transformed into *E. coli* BL21DE3-C43 cells (www.lucigen.com) and cultures were grown at 37°C and 200 rpm in 50 mL Terrific Broth (TB) medium to an OD_600_ of ~0.5-0.6 before cooling to 16°C, and induction with 1 mM isopropyl-β-D-1-thiogalacto-pyranoside (IPTG) and addition of 40 mM sodium pyruvate, 1 mM MgCl_2_, 5 mg L^-1^ riboflavin and 75 mg L^-1^ 5-aminolevulinic acid. After 72 h incubation, metabolites were extracted with hexane and air-dried for GC-MS analysis on an Agilent 7890B GC interfaced with a 5977 Extractor XL MS Detector at 70 eV and 1.2 mL min^-1^ He flow, using an Agilent DB-XLB column (30 m, 250 μm i.d., 0.25 μm film) and the following GC parameters: 50°C for 3 min, 15°C min^-1^ to 300°C, hold 3 min with pulsed splitless injection at 250°C. MS data from 40-400 mass-to-charge ratio (*m/z*) were collected after a 13 min solvent delay. Metabolite quantification (n=3) was based on normalization to the internal standard 1-eicosene (www.sigmaaldrich.com) and OD_600_ at time of induction of protein expression.

### NMR analysis

For NMR analysis, ≥1 mg of diterpene products was enzymatically produced as outlined above and purified by silica chromatography and semi-preparative HPLC as previously described (Murphy et al., 2019). Purified compounds were dissolved in deuterated chloroform (CDCl_3_; www.sigmaaldrich.com) containing 0.03% (v/v) tetramethylsilane (TMS). NMR 1D (^1^H and ^13^C) and 2D (HSQC, COSY, HMBC and NOESY) spectra were acquired on a Bruker Avance III 800 MHz spectrometer (www.bruker.com) equipped with a 5 mm CPTCI cryoprobe using Bruker TopSpin 3.2 software and analyzed with MestReNova 11.0.2 software (https://mestrelab.com/). Chemical shifts were calibrated against known chloroform (^1^H 7.26 and ^13^C 77.0 ppm) signals.

### Homology modeling and molecular docking

Homology models CYP71Z25-29 were generated with RosettaCM (Song et al., 2013) using the crystal structure of *Salvia miltiorrhiza* CYP76AH1 (Gu et al., 2019) (PDB-ID: 5YM3) as a template, and the lowest energy models were selected for docking with RosettaDock (Meiler and Baker, 2006; Davis and Baker, 2009). Input native protein structures almost invariably have regions that score poorly with force fields due to energetic strain, and minimization protocols commonly lead to increased deviation from original wild-type structure representing stable proteins. To mitigate these limitations cycles of minimization with combined backbone/sidechain restraints that are Pareto-optimal with respect to RMSD to the native structure and energetic strain reduction were used to relax the template protein structure (Nivón et al., 2013). The fulllength sequences for all targets and templates were aligned using PROMALS3D (Pei and Grishin, 2014). Each target sequence was respectively threaded onto each template and threaded partial models aligned in a single global frame. Full-chain models were then generated by Monte Carlo sampling guided by the Rosetta low-resolution energy function supplemented with distance restraints from the template structures and a penalty for separation in space of residues adjacent in the sequence. Structures were built using a Rosetta ‘‘fold tree” (Das and Baker, 2008). The global position of each segment was represented in Cartesian space, whereas the residue backbone and side-chain conformation in each segment were represented in torsion space. Using the aligned target and template sequences, evolutionary constraints were calculated and used for modeling (Thompson and Baker, 2011). A total of 500 homology models were generated for each variant, where the model with the lowest overall protein score (“total score”) was utilized for docking each substrate. First, the heme co-factor was docked into the model, followed by docking of the substrate variations [i.e., three possible C15 and C16 substitution arrangements for all five tested substrates: hydroxy (C15=OH, C16=H), pyrophosphate (C15=OPP, C16=H) and dihydroxy (C15=OH, C16=OH)] into the heme-model complex. The reactive carbon was heavily weighted within Rosetta to be constrained to a distance of 2±1 Å. Each docking simulation generated 1,000 docked poses that were filtered by “high” constraint (CST) scores, subsequently by total score (Sc) for the lowest 25%, and by interaction energy (IE) for the remaining lowest 25%.

### Site-directed mutagenesis

Select point mutants were generated using whole-plasmid PCR amplification with sitespecific sense and anti-sense oligonucleotides (Table S3) and Phusion HF Master Mix polymerase (www.neb.com). *Dpn1* treatment was applied to remove template plasmids before transformation into DH5α cells for plasmid propagation. All mutants were sequence verified and functionally characterized using *E. coli* co-expression assays as described above.

### *In vivo* feeding study

For substrate feeding assays, the constructs pETDuet-1:*Zm*CPR2/CYP71Z25 and pETDuet-1:*Zm*CPR2/CYP71Z29 were individually expressed in *E. coli* as described above. Expression of the base plasmid pETDuet-1:*Zm*CPR2 was used as a control. At the time of IPTG induction of protein expression, 10 μM of the diterpene alcohol substrates, **15** or **16**, dissolved in 1:1 (v/v) DMSO:MeOH were added to the culture, followed by incubation and metabolite analysis as outlined above.

### Metabolite profiling of tissue extracts

Frozen tissue (~500 mg) was ground to a fine powder and metabolites were extracted in hexane:ethyl acetate (Hex:EA; 80:20) with shaking overnight at 200 RPM at 12°C. Root extracts were air-dried, re-dissolved in Hex:EA, and concentrated to a volume of 1 mL for GC-MS analysis as outlined above. Leaf extracts were treated in the same manner with the addition of partially purifying extracts over a mock silica column as previously described prior to analysis by GC-MS (Muchlinski et al., 2019). Relative metabolite quantification (n=3) was based on normalization of the analytes to the internal standard (1-eicosene; www.sigmaaldrich.com) and gDW.

### Transcriptome profiling of drought-elicited switchgrass tissues

Switchgrass plants (var. Alamo AP13) were propagated from tillers to maintain low genetic variation. Plants were cultivated in 9.5 L pots in a randomized block design under greenhouse conditions to reproductive stage R1, with a 16hr light/8hr dark photoperiod and ~22/17°C day/night temperature prior to drought treatment. Drought stress treatment was applied by withholding water for four weeks, compared to control plants receiving daily drip irrigation with nutrient solution. Relative soil water content was measured weekly using a HydroSense II probe (Campbell Scientific). Leaf and root tissue of droughted and control plants (n=6) were collected weekly and flash-frozen in liquid nitrogen for later processing. Total RNA was extracted from 100 mg of tissue using a Monarch^®^ Total RNA Miniprep Kit (www.neb.com) and treated with *DNase I* for genomic DNA removal. Preparation of cDNA libraries and transcriptome sequencing was performed by Novogene Co. Ltd. (https://en.novogene.com). In brief, following RNA integrity analysis and quantitation, cDNA libraries were generated using a NEBNext^®^Ultra™RNA Library Prep Kit (www.neb.com) and sequenced on an Illumina Novaseq 6000 sequencing platform generating 40-80 million 150 bp paired-end reads per sample. Raw reads were processed using FastQC (http://www.bioinformatics.babraham.ac.uk/projects/fastqc/), and high-quality reads were aligned to the reference genome (*P. virgatum* var. Alamo AP13 v5.1) using HISAT2. Heatmaps were generated using the ‘pheatmap’ package in R (cran-project.org, version 3.6.3).

### Phylogenetic analysis

A maximum likelihood phylogenetic tree was generated using the PhyML (http://www.atgc-montpellier.fr/phyml/) server with four rate substitution categories, LG substitution model, BIONJ starting tree and 500 bootstrap repetitions (Guindon et al., 2010).

### Accession numbers

Nucleotide sequences of P450 genes and enzymes characterized in this study are available on Phytozome (https://phytozome.jgi.doe.gov): *CYP71Z25* (*Pavir.1KG341400*), *CYP71Z26* (*Pavir.1KG382300*), *CYP71Z27* (*Pavir.1KG382400*), *CYP71Z28* (*Pavir.1NG304500*), *CYP71Z29* (*Pavir.1NG309700*). Gene identifiers based on *P. virgatum* (var. Alamo AP13) genome (version v5.1). The RNA-seq data have been submitted to the Sequence Read Archive (SRA), accession no. PRJNA644234.

## Abbreviations

diTPS: diterpene synthase
P450: cytochrome P450 monooxygenase
CPS: copalyl diphosphate synthase
KSL: kaurene synthase-like
Pv: *Panicum virgatum*
GGPP: geranylgeranyl pyrophosphate
CPP: copalyl pyrophosphate
LPP: labda-13-en-8-ol pyrophosphate
CLPP: clerodienyl pyrophosphate
GC-MS: gas chromatography-mass spectrometry
HPLC: high-performance liquid chromatography
NMR: nuclear magnetic resonance
IE: interaction energy
REU: Rosetta energy units
SWC: soil water content.

## Acknowledgements

We gratefully acknowledge the US Department of Energy Joint Genome Institute (DOE JGI) and collaborators for prepublication access to the *Panicum virgatum* v1.1 and v4.1 genome sequence data produced by DOE JGI. We further thank Dr. Reuben Peters (Iowa State University, USA) for providing the pIRS and pGGxC constructs, the UC Davis Genome Center Bioinformatics Core for providing high-performance computing resources, and Dr. Malay Saha (Nobel Research Institute, USA) for providing tillers of *P. virgatum* var. Alamo AP13.

## Author Contributions

P.Z. conceived the original research and oversaw data analysis; A.M. and M.J. performed most experiments and data analyses; K.T. conducted plant drought stress experiments, metabolite profiling of plant tissues, and transcriptome analysis; J.S.F. and J.S. performed protein modeling and molecular docking studies; K.A.P. performed NMR structural analyses; L.C., D.D and Y.C. assisted with site-directed mutagenesis studies; J.T.L. performed gene synteny studies; A.M., M.J. and P.Z. wrote the article with contributions from all authors. All authors have read and approved the manuscript.

## Supplementary Information

**Fig. S1:** Identification of CYP71Z25-29 paralogs.

**Fig. S2:** Diterpenoid-metabolic pathways and compounds relevant to this study.

**Fig. S3:** Functional characterization of switchgrass CYP71Z25-29.

**Fig S4:** NMR analysis of *syn*-15,16-epoxy-8(17),13(16),14-triene (17).

**Fig S5:** NMR analysis of 15,16-epoxy-8,13(16),14-triene (18).

**Fig. S6:** NMR analysis of *cis-trans*-15,16-epoxy-cleroda-3,13(16),14-triene (19).

**Fig. S7:** Functional characterization of switchgrass P450s.

**Fig. S8:** NMR analysis of (+)-15,16-epoxy-8(17),13(16),14-triene (23).

**Fig. S9:** Substrate specificity of switchgrass CYP71Z25-29.

**Fig. S10:** Structural analysis of CYP71Z25-29.

**Fig. S11:** *In planta* accumulation of furanoditerpenoids in drought-treated switchgrass roots.

**Table S1:** Average and standard deviation for each interaction energy (IE) value of filtered docking poses for each P450-substrate combination.

**Table S2:** Synthetic genes used in this study.

**Table S3:** Oligonucleotides used in this study.

**Table S4:** Abbreviations and accession numbers for proteins used for phylogenetic studies.

